# When two mindfulnesses meet

**DOI:** 10.1101/728584

**Authors:** Louis Lakatos, Josée Turcotte, Bruce Oddson

## Abstract

The study of mindfulness proceeds from a number of perspectives. Two of the best-known academic conceptualizations of mindfulness are those identified with Kabat-Zinn and Langer. These conceptions, meditative and socio-cognitive, have been built from different foundations and have been argued to be quite distinct. However, Hart, Ivtzan and Hart ^1^ suggested that self-regulation of attention is a mediator between the two. To put this hypothesis to a test, a convenience sample of participants (n = 208) were asked to complete the Five Facet Mindfulness Questionnaire (FFMQ), Langer Mindfulness Scale (LMS), and the Self-Regulation Scale (SRS), a measure of the self-regulation of attention. These three dispositional measures were shown to be correlated. Self-regulation passes a statistical test for partial mediation of the relationship between the two measures of mindfulness. This suggests that reliance on the capacity to regulate attention in pursuit of a goal is shared by these two approaches to mindfulness. However, there is no clear conceptual basis for mediation in either particular direction. Further, the correlation between the LMS and FFMQ is highest for those with the highest SRS scores; we discuss the implications for conceptual distinctions within mindfulness.

## Introduction

In the last 35 years, there has been a broad and increasing interest in mindfulness (for examples, see ^2-8^). This interest is both academic and practical, as practices which facilitate being mindful have been linked with improvements in physical and psychological well-being including happiness, emotional regulation, neuroendocrine function and control of attention ^2, 5, 7-11^. The variety in both theory and practice has led to some conceptual challenges ^12^.

Within the western academic world, there are two main approaches to the definition of mindfulness: one, popularized by Jon Kabat-Zinn ^13^ and his colleagues, as a meditative disposition, and the other, promoted by Ellen Langer ^14^ and her associates, defined more as a socio-cognitive stance. Kabat-Zinn’s definition, broadly in line with eastern traditions, takes mindfulness as “The awareness that emerges through paying attention on purpose, in the present moment, and non-judgmentally to the unfolding of experience moment by moment” (^5^, p. 145). Being mindful according to the meditative tradition is primarily promoted by the practice of meditation, with the state defined by a combination of intention, attention, and attitude ^8^. In Langer’s view, being mindful is an orientation to being attentive no matter the context. Her conceptualization of mindfulness emphasizes being mindful of the novelty in information, the context in which it is embedded and the multitude of perspectives inherent to every situation. People are encouraged to develop cognitive flexibility and curiosity to each situation they encounter, thereby increasing their engagement and consequently their awareness. The two views on mindfulness lead to distinct types of practices, research interests, and measures ^7^.

Kabat-Zinn’s definition of mindfulness gives a central role to attention. Meditation can be viewed as the activity of holding attention in such as way that the mind does not wander – with the quality of the experience being determined by one’s success in doing so. In watching the breath, one typical mindfulness practice, the attention is directed toward the sensations elicited by the breath. The nature of attention is such that this will require an ongoing effort to maintain this direction, against a tendency (for beginners) to wander away. Self-regulation of attention is needed to monitor the state of attention, detect wandering, and bringing it back to the breath.

This view is by no means idiosyncratic. In a review, Khoury et al. ^15^ found that eight of the ten western academic definitions of mindfulness made attention part of their foundation, as do some of the Buddhist sources. The remaining definitions referred to mental engagement and awareness, which in the terms of cognitive psychology amounts to the same thing. Thus, mindfulness in the moment is described in attentional terms directly or in terms of the mechanism which guide awareness, and failures to be mindful result in loss of intended focus - mind wandering. Morrison and Jha ^16^ present data supporting the link between attention and mind wandering. They go further to suggest that practices of mindfulness (both within a session and over longer periods of time) strengthen specific aspects of attention (and working memory). Generally, dispositional mindfulness measures are associated to measures of attention ^17^ which is also consistent with the idea that the control of attention is central both to meditation and its aftereffects ^18^.

One of the most popular measures of meditative mindfulness is the Five Facet Mindfulness Questionnaire (FFMQ ^19^). Rather than describing the state of mindfulness in the moment, the FFMQ measures a disposition, a tendency to be mindful in general in everyday life. Accordingly, its scores need to be assumed to reflect a broad capacity for the direction of attention. The FFMQ was created to assess the five key mechanisms supporting mindfulness: observing (noticing internal and external stimuli), describing (ability to assign verbal labels to experiences), acting with awareness (paying attention to the present moment), nonjudging of inner experience (inhibit evaluation of thoughts and emotions), and nonreactivity to inner experience (ability to allow stream of consciousness to flow freely without restraint or evaluation). These factors do not all relate to attention as a mechanism in the same way (for a discussion of the factors see Rau and Williams ^20^). Some of the scores (e.g. observing and acting with awareness) lend themselves to description in attentional terms. The status of the other subscales is less clear, as non-judgment and non-reactivity are traditionally considered fruits of mindfulness practice as much as signs of successful direction of attention.

Langer and her colleagues developed their concept of socio-cognitive mindfulness in the 1970s. In this concept, the primary theoretical distinction is between mindlessness – acting and thinking largely from learned experience – and mindfulness in which attention is kept on variation and interest in the present. This distinguishes mindfulness from a description or byproduct of meditative practices. Rather, in this school of thought mindfulness was conceived as a mental mode grounded in a person’s disposition (^14^; see also ^21^, p.22)). Langer’s conception encourages us to be mindful of the novelty in information. It is defined on cognitive flexibility and curiosity in each situation we encounter, thereby increasing our engagement and consequently our awareness ^7,22^.

To achieve mindfulness in this sense requires an active engagement in the present and an effort to actively draw novel distinctions in the context in which it is embedded and the multitude of perspectives inherent to every situation ^7^. Through practice, this eventually becomes part of a person’s disposition ^7, 23^. The involvement of self-regulation of attention is not explicitly stated as a central mechanism in the conception of socio-cognitive mindfulness but the necessity of being open to novelty, engaged and actively drawing novel distinctions emphasizes its importance ^7^. To avoid mindlessness requires both a willingness and a capacity to make an effort with our attention. We would not only need less automatic and habitual responses but also more self-control “to over rides prepotent response” (^24^, p. 884) in order to focus one’s awareness on external stimuli in a novel way. In that sense, it could be considered self-regulation of attention.

The Langer Mindfulness Scale (LMS ^25^) was constructed to capture this. Conceptually, it measures a disposition consisting of four dimensions: 1) engagement - being aware of changes that take place in the environment; 2) seeking novelty - having an open and curious orientation to one’s environment; 3) novelty producing - the capacity to construct new meanings or experiences; 4) flexibility - the tendency to view experiences from multiple perspectives and to adjust one’s behavior accordingly. The latter dimension has been excluded from the LMS14 ^26^. Of these dimensions, engagement has the most obvious link with attention. It is hypothesized that engagement reflects a continuous effort of attention to the present circumstance - and a comparative reduction of automaticity in thought ^1,26^.

A recent paper by Hart, Ivtzan, and Hart ^1^ contrasting the meditative and socio-cognitive views suggested that, despite major apparent differences, they may rest on a common mechanism – self-regulation of attention. Briefly, their argument is that since the meditative view is directly defined on attention and the socio-cognitive view of mindfulness equally rests on an ongoing effort to direct attention in a particular way, it makes sense that self-regulation of attention would be an important common factor between the two.

According to Baumeister, Vohs and Tice ^27^(p.351), “Self-control [conscious self-regulation] refers to the capacity for altering one’s own responses, especially to bring them into line with standards such as ideals, values, morals, and social expectations, and to support the pursuit of long-term goals. … Self-control enables a person to restrain or override one response, thereby making a different response possible”. Since it is effortful, self-control seems to have a limited capacity in the short term. But it may also improve through training, and become more automatic ^27^. The Self-Regulation Scale (SRS ^28^) was created to capture the capacity to regulate attention and emotions in the service of goal directed actions. The questions (e.g. “I can concentrate on one activity for a long time, if necessary”) specifically address self-perceived capacity to direct attention as needed. Again, the definitions of mindfulness describe actions of attention, giving the self-regulation of attention a potential role.

### Present Study

The socio-cognitive model championed by Langer places its focus on the creative application of mindfulness as opposed to automaticity to promote novelty and engagement. The meditative view, often associated with Kabat-Zinn, indicates that mindfulness involves (a) self-regulation of one’s awareness, (b) directing one’s attention to internal and external stimuli, (c) introspection and metacognitive awareness of one’s thoughts processes, and (d) adopting a nonjudgmental attitude ^3^. Hart et al. ^1^ contrasted these views and suggested that self-regulation of attention may be an important point in common between these two views. In the present work we investigate this claim empirically.

Previous literature has considered the possible correlation between the LMS and FFMQ - the measures closely associated with the two schools of thought. Trend, Park, Bercovitz and Chapman ^29^ found a correlation (r = .49) between them using a large sample of the general population (N=484). Siegling and Petrides ^30^ found that their total scores were weakly correlated in two samples (r = .33 and r = .36). They also found that the LMS related to personality factors in a notably different way from the FFMQ and other meditative mindfulness measures. Other studies have found similar correlations (e.g. Pirson et al, 2018 :^26^) with some studies finding no correlation beyond what is expected by chance (Pirson, Langer, Bodner and Zilcha, 2012, as cited by Siegling and Petrides, ^31^).

In principle, there are many reasons why the correlations between the LMS and FFMQ might be positive in some cases and absent in others. They do measure different constructs. However, there are also some measurement considerations. A limited range of scores will tend to attenuate correlations, as will measurement error. Accordingly, samples in which mindfulness is generally low may have weaker correlations than samples which cover more of the possible range. We will take up this point in the discussion. Both scales have acceptable internal consistency, but neither was intended to produce a unidimensional score. However, from a conceptual point of view, there are two reasons to expect some correlation. The first is that Hart et al. ^1^ have correctly identified attention as a common foundation. The second is that mindfulness, meditative or socio-cognitive, has cognitive consequences.

In the present study, the main hypothesis is that self-regulation of attention mediates the relationship between meditative (as measured by the FFMQ) and socio-cognitive (measured by the LMS) mindfulness. In other words, that the capacity for self-regulation of attention would explain the relation between the two type of dispositional mindfulness. However, given the previous literature and the analysis by Hart et al. ^1^, we also consider the relationship between the subscales.

## Materials and methods

### Measures

#### FFMQ

The FFMQ scale is a 39-item comprehensive mindfulness scale based on meditative mindfulness, and it measures five facets of the construct; observing, describing, acting with awareness, nonjudgment of inner experience, non-reactivity to experience ^19^. The FFMQ was derived from confirmatory factor analyses of previous scales. The FFMQ has a Cronbach’s alpha coefficient of 0.90, indicating good reliability. Item response categories range from 1 ‘Never or very rarely true’ to 5 ‘Very often or always true’.

#### LMS

The LMS is a 21-item Likert scale based on socio-cognitive mindfulness, and it includes the following subscales: flexibility, novelty producing, novelty seeking, engagement (Langer, 2004). The LMS had a Cronbach’s alpha coefficient of 0.85, indicating good reliability. Item response categories from 1 ‘strongly disagree’ to 7 ‘strongly agree’.

#### SRS

The SRS is a 10-item scale designed to measure self-regulation capabilities; specifically, self-regulation of attention in the pursuit of a goal ^28^. The coefficient of internal consistency (Cronbach’s α) was .76. Item response categories range from 1 ‘not all true’ to 4 ‘completely true’.

#### Socio-demographic questionnaire

Questions regarding the participant’s age, sex, education, meditation experience were asked before the main scales were presented.

### Participants and Recruitment

The study was approved by the Research Ethics Board (REB) at Laurentian University. Participants were recruited using three methods: 1) through undergraduate university classes, 2) with snowball sampling technique, and 3) through online advertising. This was done to maximize recruitment and allow the recruitment of a variety of participants in terms of age, meditation experience, etc. The first method was in class student recruitment, where undergraduates were offered the chance to participate in this research in exchange for bonus marks in academic courses. The second involved the snowball technique which functions like chain referral. Participants were asked to pass a survey along to family and friends who might be interested in mindfulness. The third method involved sending the invitation to various mindfulness centers where several managers were contacted and requesting that they share the survey to interested clients and parties. Those who completed the survey were also given an option to be entered into a draw for a $50 Amazon gift card by sharing the survey link over social media in exchange for points which would determine a winner based on the greatest number of shares.

In total, 261 participants were recruited and started the survey; 208 of these participants answered all questions and were included in the analysis. Amongst the participants, 176 identified themselves as females (33 males). The mean age was 23.84 (SD = 10.20). One hundred fifty-four people reported no meditation experience with 55 people reporting previous meditation experience of some kind (yoga, sitting meditation, etc.). Please see the demographic characteristics in Table 1 for more information.

**Table 1.**
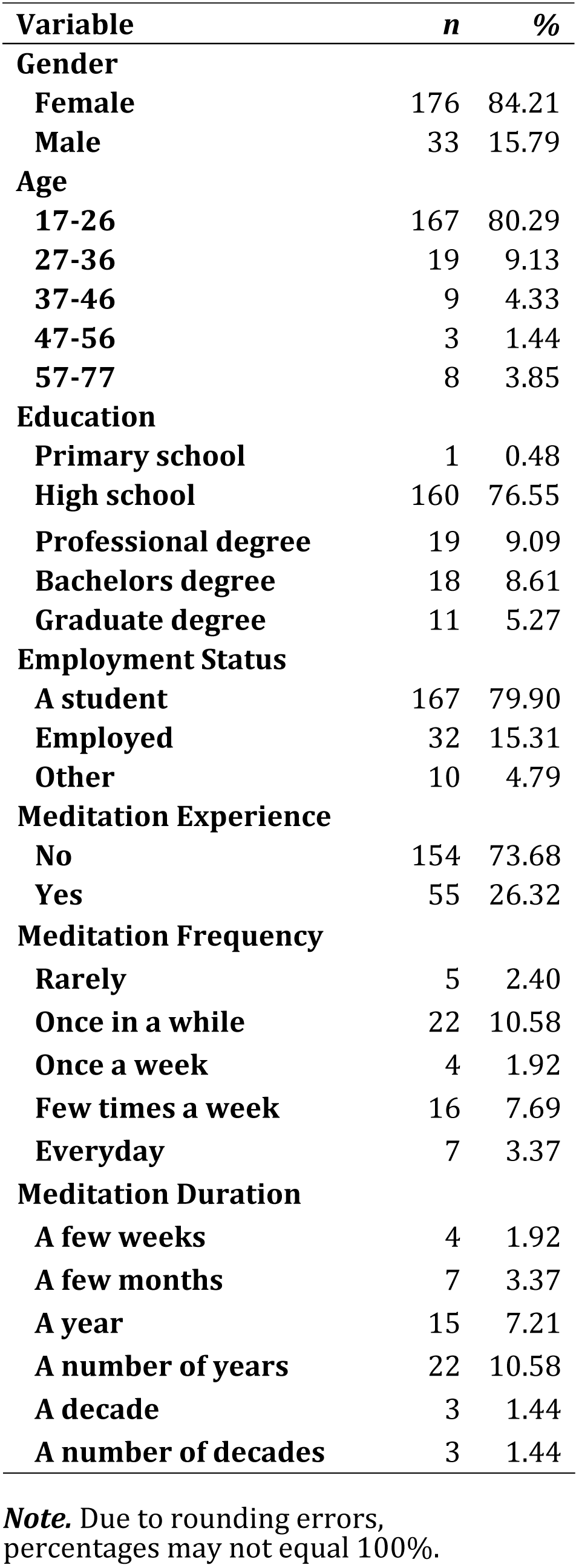
Demographic Characteristics.

| **Variable** | ***n*** | **%** |
| --- | --- | --- |
| **Gender** |  |  |
| **Female** | 176 | 84.21 |
| **Male** | 33 | 15.79 |
| **Age** |  |  |
| **17-26** | 167 | 80.29 |
| **27-36** | 19 | 9.13 |
| **37-46** | 9 | 4.33 |
| **47-56** | 3 | 1.44 |
| **57-77** | 8 | 3.85 |
| **Education** |  |  |
| **Primary school** | 1 | 0.48 |
| **High school** | 160 | 76.55 |
| **Professional degree** | 19 | 9.09 |
| **Bachelors degree** | 18 | 8.61 |
| **Graduate degree** | 11 | 5.27 |
| **Employment Status** |  |  |
| **A student** | 167 | 79.90 |
| **Employed** | 32 | 15.31 |
| **Other** | 10 | 4.79 |
| **Meditation Experience** |  |  |
| **No** | 154 | 73.68 |
| **Yes** | 55 | 26.32 |
| **Meditation Frequency** |  |  |
| **Rarely** | 5 | 2.40 |
| **Once in a while** | 22 | 10.58 |
| **Once a week** | 4 | 1.92 |
| **Few times a week** | 16 | 7.69 |
| **Everyday** | 7 | 3.37 |
| **Meditation Duration** |  |  |
| **A few weeks** | 4 | 1.92 |
| **A few months** | 7 | 3.37 |
| **A year** | 15 | 7.21 |
| **A number of years** | 22 | 10.58 |
| **A decade** | 3 | 1.44 |
| **A number of decades** | 3 | 1.44 |

**Note.** Due to rounding errors, percentages may not equal 100%.

### Procedure

All participants completed the survey through the Redcap system which is an online survey creation website that is commonly used in university research. First, participants read a notification statement that explained the risks and benefits of participating and emphasized their rights to skip any answer or withdraw from the study at any time. They provided consent to participate by clicking “Next” in the survey and reported demographic information consisting of age, gender, education, marital status and employment. Additional questions asked about meditation experience (type, duration and reasons for practice). The three main scales of the FFMQ, LMS, and SRS were presented next and concluded the survey process. The entire survey required approximately 1 hour to complete. All this

## Data Analysis

The correlations between the FFMQ, LMS, SRS, demographic variables and relevant subscales were assessed using Pearson correlations. Mediation by SRS of the relationship between the LMS and FFMQ was assessed using the Sobel-Goodman test; as were tests of mediation involving their subtests. Finally, a finite mixture model was generated treating SRS as a latent class to investigate the role of SRS in their relationship. All analyses were conducted using STATA 15.1.

## Results

The Table 2 presents the summary statistics (mean, standard deviations and range of scores) of the FFMQ, LMS and SRS. Mindfulness, as measured by the FFMQ total score, is correlated with mindfulness, as measured by the LMS total score, r = .52, p < .01, 95% CI [.42 - .61]. Thus, people who are high in meditative mindfulness tend to be high in socio-cognitive mindfulness. A similar result is found if the LMS total score is recalculated using the 14 items of the revised version (LMS14; Pirson, et al., 2018), r = .46, p < .01, 95% CI [.34 - .56]. Many of the subscales of the FFMQ are also correlated with those of the LMS (please see Table 3); the exception being the non-judgment subscale.

**Table 2.** Summary Statistics and the normality of the FFMQ, LMS and the SRS.

| Variable | *M* | *SD* | *Range* |
| --- | --- | --- | --- |
| **FFMQ** | 122.31 | 17.93 | 64-179 |
| **LMS** | 107.40 | 14.12 | 70-143 |
| **SRS** | 27.28 | 5.16 | 13-40 |

**Legend.** FFMQ: Five-Facet Mindfulness Questionnaire; LMS: Langer Mindfulness Scale; SRS: Self-Regulation Scale.

**Table 3.** Correlations and 95% Confidence Intervals between the FFMQ and LMS.

| FFMQ | LMS |  |  |  |
| --- | --- | --- | --- | --- |
|  | Flexibility | Nov. Seeking | Nov. Producing | Engagement |
| Observing | $r = .35$ (.23 - .47) | $r = .40$ (.28 - .51) | $r = .45$ (.33 - .55) | $r = .38$ (.26 - .49) |
| Describing | $r = .26$ (.13 - .38) | $r = .25$ (.12 - .37) | $r = .31$ (.18 - .43) | $r = .36$ (.24 - .48) |
| Act. w Awareness | $r = .18$ (.05 - .31) | $r = .19$ (.06 - .32) | $r = .24$ (.11 - .37) | $r = .39$ (.27 - .50) |
| Non-Judgement | $r = .14$ (.00 - .27) | $r = .07$ (-.07 - .20) | $r = .04$ (-.10 - .17) | $r = .22$ (.09 - .35) |
| Non-Reactivity | $r = .42$ (.30 - .52) | $r = .27$ (.14 - .40) | $r = .34$ (.21 - .45) | $r = .32$ (.19 - .44) |

**Legend:** Act. w Awareness : Acting with Awareness, Nov. Seeking : Novelty Seeking, Nov. Producing : Novelty Producing

The FFMQ total score correlates with the SRS, r = .66, p < .01, 95% CI [.58 - .73], as do the subscale scores, with the highest correlation going to acting with awareness, r= .63, 95% CI [.54 - .70], and the lowest with observing, r= - .26, 95% CI [.12 --.38]. These correlations reveal that people who are high in any aspect of meditative mindfulness tend to have a high score in the self-regulation of attention. The same is true for the LMS, r = .45, p < .01, 95% CI [.34 - .55], and its subscales (all rs > .3) indicating that people who are high in any aspects of socio-cognitive mindfulness also tend to be high in self-regulation of attention.

Using the Sobel-Goodman procedure, it is estimated that the SRS explains about 46% of the correlation between the FFMQ and the LMS, when the FFMQ is the dependent variable and the LMS is the independent variable. It suggests that SRS is a partial mediator of their relationship, z = 5.76, p < .01. Both the direct and indirect effects are significant, ps < .01. Due to the asymmetric relationship, when the FFMQ is considered the independent and the LMS is become the dependent variable, the mediation effect is estimated to be weaker, about 22%, but still significant, p <.01.

Partial mediation suggests that SRS plays a role in mindfulness but is not the only point of contact between the LMS and FMQ. To explore the relationship further, a latent class model was created. We tested whether or not the correlation between the FMQ and LMS was stable over the entire sample. Membership in one of two latent groups were identified by SRS score. This model showed that the SRS score could make a clear distinction between groups, z = 3.18, p = .002, such that the correlation between LMS and FMQ was low, r = .26, p < .001 when SRS scores are not high, and the correlation was high, r = .64, p < .001, for a group centered on the highest SRS quintile (see Table 4).

**Table 4.**
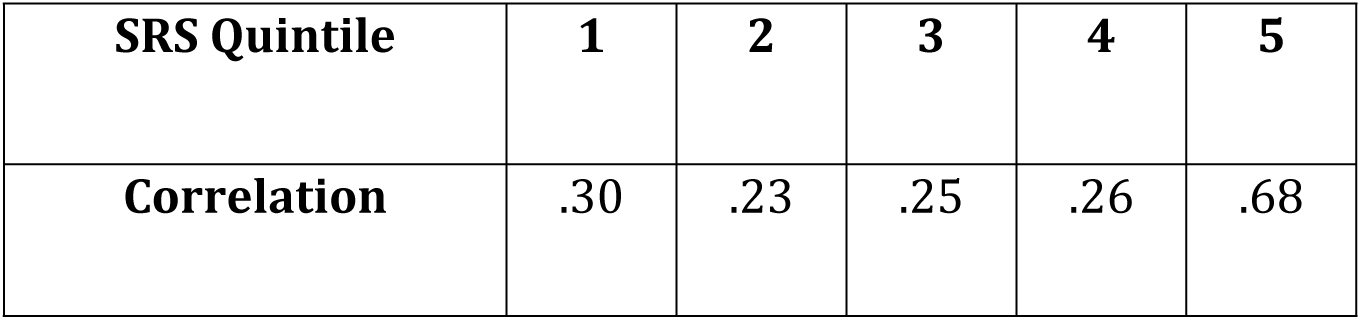
Correlation between LMS and FMQ by SRS Quintile.

| SRS Quintile | 1 | 2 | 3 | 4 | 5 |
| --- | --- | --- | --- | --- | --- |
| Correlation | .30 | .23 | .25 | .26 | .68 |

## Discussion

There is a significant and growing literature about the conceptual and empirical relationships between the approaches to mindfulness we have labeled meditative and socio-cognitive. These approaches have major differences in their conceptual foundations, and there are good reasons to suppose that these differences would be reflected in their measures. However, there are also several points of connection between meditative and socio-cognitive mindfulness. Meditative mindfulness is intended to capture a disposition for holding the mind in states consistent with meditation - concentrated, with awareness enhanced and judgment suspended. Socio-cognitive mindfulness is also defined as a disposition - remaining engaged with a flexibility and depth that supports high level cognition and creativity.

Hart et al. ^1^ analyzed these two conceptions and created an argument that the two conceptions should be linked by the capacity to control attention. In fact, many of the conceptual reviews of mindfulness give a leading role to attention, with the dispositional marker for attentional capacity being self-regulation ^10^. We assert that self-regulation of attention is an appropriate potential mediator to test. Hart et al. (2013) make the additional prediction that greatest overlap between the tests should rest on the observing and acting with awareness FFMQ subscales and the engagement LMS subscale.

In the present study, the FFMQ, a measure developed within the meditative approach to mindfulness, and the LMS, a measure developed by the leading exponent of the socio-cognitive approach, were correlated. Our data provide some support to the Hart et al.^1^ predictions, but also opens additional questions. We find that the correlation between the FFMQ and LMS is partially mediated by scores on the SRS. These results support the claim that self-regulation of attention can be identified as a central aspect of both approaches. However, there are a few complications in interpreting this finding. One of the first is that the Sobel-Goodman test presumes a direction of causality. There is no reason to suppose a particular direction of influence between these two measures; however, the size of the mediation effect does depend on direction (due to the asymmetric correlations with the SRS). In either case, there is a sizeable direction correlation between these measures even once the indirect pathway involving attention has been considered. Testing the regression interaction term, as per moderation analysis, leads to the same conclusion. This suggests that the position of Hart et al. ^1^ that self-regulation of attention is a central aspect of both is supported, but that this is not a complete description.

A measurement approach to this finding would have to acknowledge that none of the measures involved are ideal. To the extent that the FFMQ and LMS tap into aspects of attention not measured by the SRS, the size of the mediation effect found should not be taken to reflect negatively on the Hart prediction. The FFMQ is made up of five subscales and the LMS has four. As presented in Table (3), most of the possible correlations are significant, with the exception of those involving non-judgment. In fact, these correlations include the observing subscale - which has been the most criticized in terms of its role within the FFMQ ^32^ and the flexibility subscale from the LMS, which has been dropped from the revised version ^26^ due to weak loadings on the factor structure of the remaining items. It is worth nothing that our results remain consistent if the FFMQ is rescored without the observing score and compared with the revised LMS14. It is also worth considering that the correlations between each of the measures are attenuated by their reliability coefficients - and the underlying relationship could then be stronger than what is observed.

From a conceptual point of view, the other possibility is that the FFMQ and LMS have more in common than has been generally supposed. There are two arguments to consider. First is that the two scales were designed as broad survey instruments. Each tap into a wide variety of experiences and self-judgments. Personality traits, their common associations with physical and psychological wellbeing, and other correlates may provide additional mediating variables. The other possibility is that, quite simply, mindfulness of the meditative sort (as with the socio-cognitive sort) has broad implications for the way we engage with the world. This last is certainly consistent with the broad holistic roots of both conceptions.

However, Hart et al.’s specific prediction about which subscales would be most correlated was not supported. The FFMQ subscales acting with awareness and observing were not clearly more correlated with the LMS subscales than were describing and non-reactivity; nor was there any pronounced tendency for correlations to engagement to be stronger than to the other LMS scales. This contradicts the specific predictions made by Hart et al. ^1^ and adds to the evidence that the connection between these tests is broader than what might be expected from the face descriptions of the questionnaires.

Whether the correlation between meditative and socio-cognitive mindfulness is indirect, through multiple mediators, or direct, reflecting a commonality of underlying disposition, the strength of the correlation found has some implications. This correlation is in line with the magnitude expected between tests of intelligence - with strongly overlapping underlying definitions but different measurement structures. In this case, not only are the measurement tools derived differently, but they have been validated with rather different experimental interventions. Could meditation practice be expected to influence socio-cognitive mindfulness? Can Langer’s brief orientation intervention affect meditative mindfulness over the long term? In the context of the present findings, the interaction between possible interventions and the relevant measures may give insight not only into the construction of these measures but also into the underlying meaning of mindfulness.

These results also bring into focus the question of population. Previous research had been inconsistent about the strength of any correlation between the FMQ and LMS. Our findings suggest that this relationship is generally weak, but can be strong for those who are high on self-regulation of attention. Researchers can expect to see different correlations depending on the population from which they take their samples.

People who regulate their attention and are sensitive to the actions of their attention may interpret the questions on the FMQ and LMS differently from those who do not. Alternately, there may be a point at which mindfulness is stable enough to have consequences - thus the population reporting relatively high scores on the SRS, FMQ and LMS may be different in a number of ways. However, the SRS is an indicator of any latent population differences. Accordingly, more research into the development of mindfulness and the roles of self-regulation of attention would be helpful. That said, the correlation found, as predicted by Hart et al. ^1^ is a first step towards reconciling the two literatures on mindfulness.

## Limitations

However valid a test, it should not be expected to perform equally well in every population to which it is administered. Other researchers have reported lower correlations between the FFMQ and LMS than was found in this study. Differences between our sample and those involved in other studies may be able to explain this. The present study used different strategies of recruitment, amongst university students as well as in the community. We may have more diversity than is typical of the literature. However, although the sample size in the present study is adequate for the estimates of correlations, the questionnaires did not gather enough detail about meditation practice, or other potentially relevant factors to be able to address possible differences. The questionnaires were administered through an online portal, and this may also affect results in ways not yet explored.

Despite these limitations, the present study is important in supporting the hypothesis that self-regulation of attention would be a common mechanism between meditative mindfulness and socio-cognitive mindfulness.

## Summary

The FFMQ and LMS, measures of mindfulness derived from two different approaches, are shown to be correlated in this study. The specific pattern of correlations found suggests that all parts of the FFMQ except the non-judgment subscale were correlated with the LMS subscales, and that engagement, the LMS subscale most closely defined to capture the role of attention, was not more correlated with FFMQ scales than were the other LMS subscales. The correlation between the FFMQ and the LMS is contingent on the SRS, a measure of self-regulation. Both the finding of partial mediation, and the results from the latent mixture model suggest that the two tests may have more in common than has been described in the past. This may mean that the tests reflect a common suite of mediating variables. It may also mean that the two conceptions reflect a common underlying whole. In an applied sense, it may be worthwhile to combine meditative and socio-cognitive interventions and to further explore whether strategies enhancing self-regulation can be used to support both.

